# Mouse-to-human modeling of microglia single-nuclei transcriptomics identifies immune signaling pathways and potential therapeutic candidates associated with Alzheimer’s disease

**DOI:** 10.1101/2025.02.07.637100

**Authors:** Alexander Bergendorf, Jee Hyun Park, Brendan K. Ball, Douglas K. Brubaker

## Abstract

Alzheimer’s disease (AD) is a progressive neurodegenerative disease characterized by memory loss and behavior change. Studies have found that dysregulation of microglial cells is pivotal to AD pathology. These mechanisms have been studied in mouse models to uncover potential therapeutic biomarkers. Despite these findings, there are limitations to the translatable biological information from mice to humans due to differences in physiology, timeline of disease, and the heterogeneity of humans. To address the inter-species discrepancies, we developed a novel implementation of the Translatable Components Regression (TransComp-R) framework, which integrated microglia single-nuclei mouse and human transcriptomics data to identify biological pathways in mice predictive of human AD. We compared model variations with sparse and traditional principal component analysis. We found that both dimensionality reduction techniques encoded similar AD disease biology on mouse principal components with limited differences in technical performance. Several mouse sparse principal components explained high amounts of variance in humans and significantly differentiated human AD from control microglial cells. Additionally, we identified FDA-approved medications that induced gene expression profiles correlated with projections of healthy human microglia on mouse principal components. Such medications included cabergoline, selumetinib, and palbociclib. This computational framework may support uncovering cross-species disease insights and candidate pharmacological solutions from single-cell datasets.

## INTRODUCTION

Microglia are resident innate immune cells actively involved in the brain’s central nervous system (1). Microglia contribute to neural homeostasis by clearing cellular debris, engaging in phagocytosis to eliminate apoptotic cells, and removing misfolded proteins (2,3). These activities are essential for promoting a healthy brain state; however, the dysregulation of these cells contributes to downstream neuroinflammation and potentially neurodegenerative disorders, such as Alzheimer’s disease (AD). AD is the most common form of dementia and is often characterized by cognitive impairment and memory loss. In the United States, AD affects more than 6.7 million people, with the number of cases escalating with the aging population (4). Key hallmarks associated with AD are the presentation of misfolded amyloid-beta (Aβ) aggregations and hyperphosphorylated tau tangle proteins in the brain (5,6).

Multiple pre-clinical studies have investigated the role of microglia in AD (7,8). Microglia have been shown to be inefficient at performing phagocytosis dependent on amyloid plaque levels in mice (9). Researchers have also reported the prolonged release of inflammatory cytokines, including TNFα and IL-1β, in aged AD mice models by microglia (10). Microglia have a role in diminishing neurons and dendritic spines in 5xFAD mice (11). In addition, Aβ promotes cognitive dysfunction by increasing the number of synapses removed by microglia through the complement system (12). Despite these connections between microglia and AD, the exact mechanisms by which altered microglia states effect AD pathology are complex to understand in humans.

A significant limitation on comparing mouse and human studies is that data across these models is not directly translatable (13,14). This limitation is often due to the differences between rodent and human physiology and the heterogeneity of the human population, which makes translating pre-clinical models to human relevance difficult. In AD studies, transgenic mouse models are prevalently used to emulate the hallmark characteristics of Aβ and tau hyperphosphorylation accumulation in the brain (15). AD mouse models also provide crucial insights into the pathological progression of the disease through controlled experimental conditions as opposed to the heterogeneity of human data, especially in post-mortem brain samples (16). Because of these differences, we must consider the results from both models to inform AD pathology. Addressing this translational challenge may help improve our understanding of AD pathology associated with microglia.

Previous groups have developed several computational methodologies to address these cross-species discrepancies and other issues with biological modeling. Deep learning models have been used to identify conserved biological mechanisms across species. For example, Nvwa leveraged a deep learning approach to predict gene expression and potential sequences cross-species (17). Also, the tool AutoTransOP used autoencoders to embed mice and human data into a shared latent space to extract cross-species insights (18). Traditional machine-learning models have also been used to perform this task. For example, Found In Translation finds the direct relationship of genes across species to identify conserved molecular features (19). However, there has been limited exploration into computational cross-species translation at a single-cell level, particularly in AD.

To address cross-species discrepancies, we developed a computational framework termed Translatable Components Regression (TransComp-R), which has performed similarly to other developed methodologies (14). TransComp-R runs principal component analysis (PCA) on a mouse dataset and projects the human data into the mouse PC space. Principal component regression is run on the resultant human PC projections to determine which PCs have significantly translatable information from a mouse model to a human dataset. This allows for elements of mouse model biology that are translatable to human biology to be identified and interrogated for human disease insights at a high resolution. This workflow is also not limited by disease-specific biology or different data formats (14,20–22).

Previously, our group applied TransComp-R to bulk RNA-seq data, yet we did not explore its application to single-cell RNA sequencing. Single-cell RNA sequencing (scRNA-seq) has deciphered distinct attributes of many types of cells amongst diseases often obscured in bulk RNA sequencing (23). A unique aspect of scRNA-seq is the inflated number of zero reads due to the immense diversity of cellular gene expression profiles, and dropout reads due to sequence technology errors (24). The resulting data can be very sparse, adding an additional layer to the complexity of the cross-species translation issue.

Here, we developed a novel implementation of TransComp-R, which extended the existing workflow for application to single-nuclei RNA-sequencing (snRNA-seq). This allows us to study cross-species gene expression at a single-cell resolution instead of a bulk-sequencing scenario, which only provides generalizable gene expression information across samples (25). We incorporated and benchmarked sparse principal component analysis (sPCA), a form of PCA which introduces sparsity into the principal components and conducted a screening analysis to determine drugs inversely associated with AD microglial features across species.

## RESULTS

### Annotation of mouse and human brain samples reveals similar cell types across species

Publicly available single-nuclei RNA sequenced data was obtained from brain hemisphere samples in WT and 5xFAD mouse transgenic lines (GSE208683) and post-mortem human AD and control brain samples from BA41/42, BA6/8, and anterior hippocampal cortex regions (GSE175814) (**Figure 1A**) (26,27). To translate RNA-sequenced information from mouse to human, we first identified 15,532 orthologous gene pairs. We filtered human subjects for cellular quality control metrics, which resulted in 10,657 AD cells (AD Subject 1 = 6,117, AD Subject 2 = 4,540) and 16,130 control cells (control subject 1 = 5,275, control subject 2 = 10,855). All samples were normalized and integrated to account for differences in batches. After integration and feature selection, we merged the samples and performed PCA for downstream clustering.

**Figure 1.**
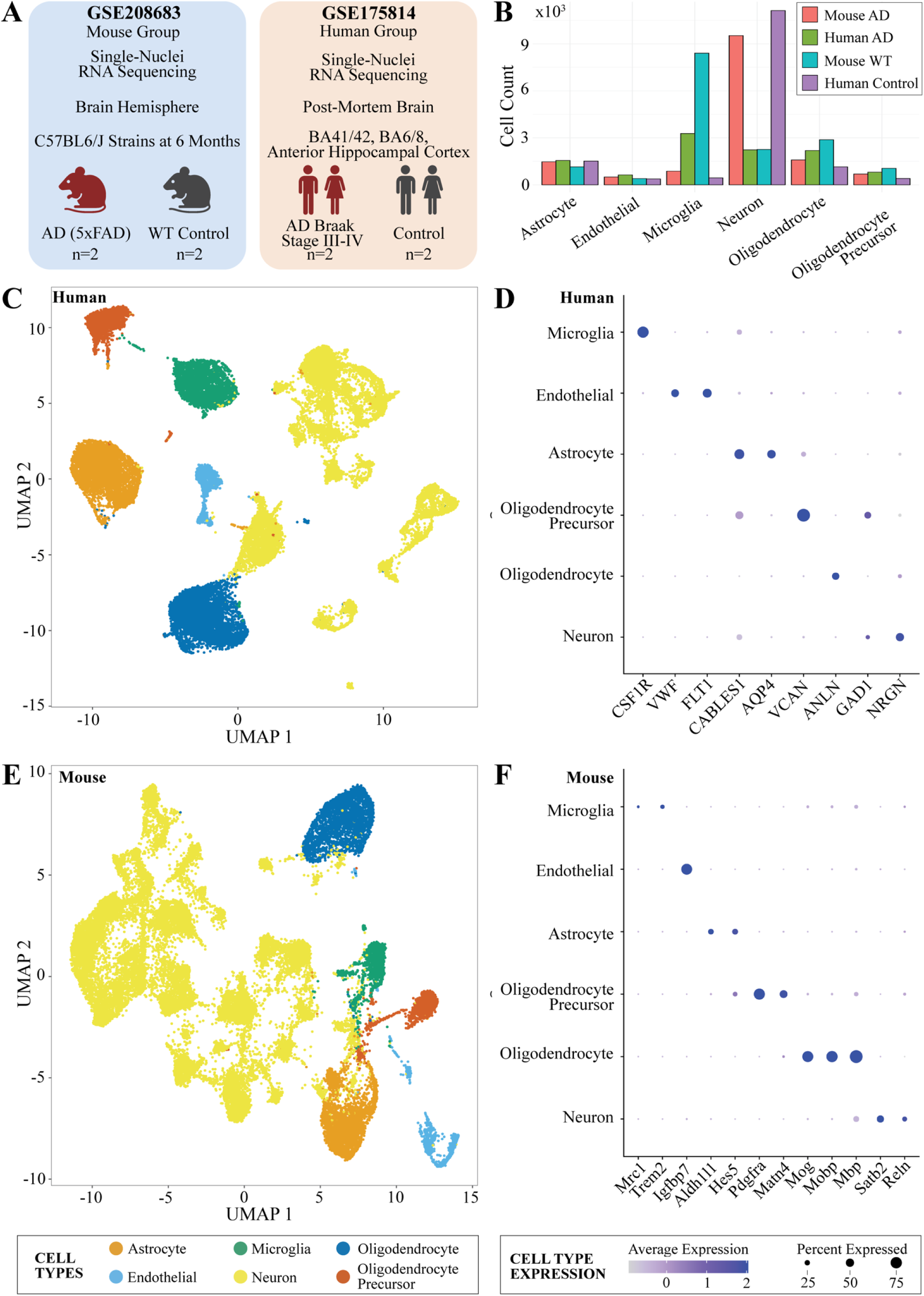
Single nuclei RNA-sequencing of human and mouse brain data. **(A)** Summary of mouse and human studies. **(B)** Cell counts for AD and control samples in studies. **(C)** UMAP of human single nuclei RNA-sequencing cells. **(D)** Marker genes of cell types in humans. **(E)** UMAP of mouse single nuclei RNA-sequencing cells. **(F)** Marker genes of cell types in mice.

From our clustering of the mouse data, 795 microglial cells pertained to AD mouse samples (5xFAD mouse 1 = 463, 5xFAD mouse 2 = 332), and 363 microglial cells came from the WT samples (WT mouse 1 = 243, WT mouse 2 = 120) (**Figure 1B**). For the human UMAP of the cells, we determined one cluster with 2,701 microglial cells, 1,553 of them corresponding to AD samples (AD subject 1 = 1,381, AD subject 2 = 172), and the remaining 1,148 microglial cells belonging to the control samples (control subject 1 = 59, control subject 2 = 1,089) **(Figure 1B)**.

We generated a KNN graph of human cells incorporating the top 30 variable PCs and a clustering resolution of 0.6. The UMAP visualization resulted in 21 different clusters from the cells (**Figure 1C**). From clustering, we identified and annotated the major cell types of the brain based on known marker genes, including microglia (*CSF1R*), endothelial including pericytes (*VWF, FLT1*), astrocytes (*CABLES1, AQP4*), oligodendrocyte precursors (*VCAN*), oligodendrocytes (*ANLN*), and neurons (*GAD1, NRGN*) (**Figure 1D**) **(Supplementary Table 1)** (28).

We also interpreted mouse cells with a KNN graph by incorporating the top 30 variable PCs and using a clustering resolution of 1.0. UMAP produced a visualization of 35 different clusters for the cells **(Figure 1E)**. From the generated UMAP of the mouse cells, we found three clusters containing 976 microglial cells. Based on the clustering, we identified and annotated the major cell types of the brain through established marker genes, including microglia *(Mrc1, Trem2)*, endothelial including pericytes *(Igfbp7)*, astrocytes *(Aldh1l1, Hes5)*, oligodendrocyte precursors *(Pdgfra, Matn4)*, oligodendrocytes *(Mog, Mobp, Mbp)*, and neurons *(Satb2, Reln)* **(Figure 1F, Supplementary Table 2)** (28–31).

### Alzheimer’s disease microglia from mice and human models shared limited differentially expressed genes

Microglia cells have several functions in the brain during neural homeostatic and disease pathology **(Figure 2A**). Some functions include the surveillance of abnormalities and foreign objects in the brain, supporting the CNS, performing phagocytosis of AD hallmarks, and releasing cytokines for pro-inflammatory or anti-inflammatory responses (32). To determine how these functions of microglia relate to AD, we investigated their expressed genes. Modeling this approach of comparison of differentially expressed genes (DEGs) across mice and humans, we ran a differential expression test using the *MAST* statistical framework. The 5xFAD and human AD samples shared only 11 orthologous DEGs (**Figure 2B**).

**Figure 2.**
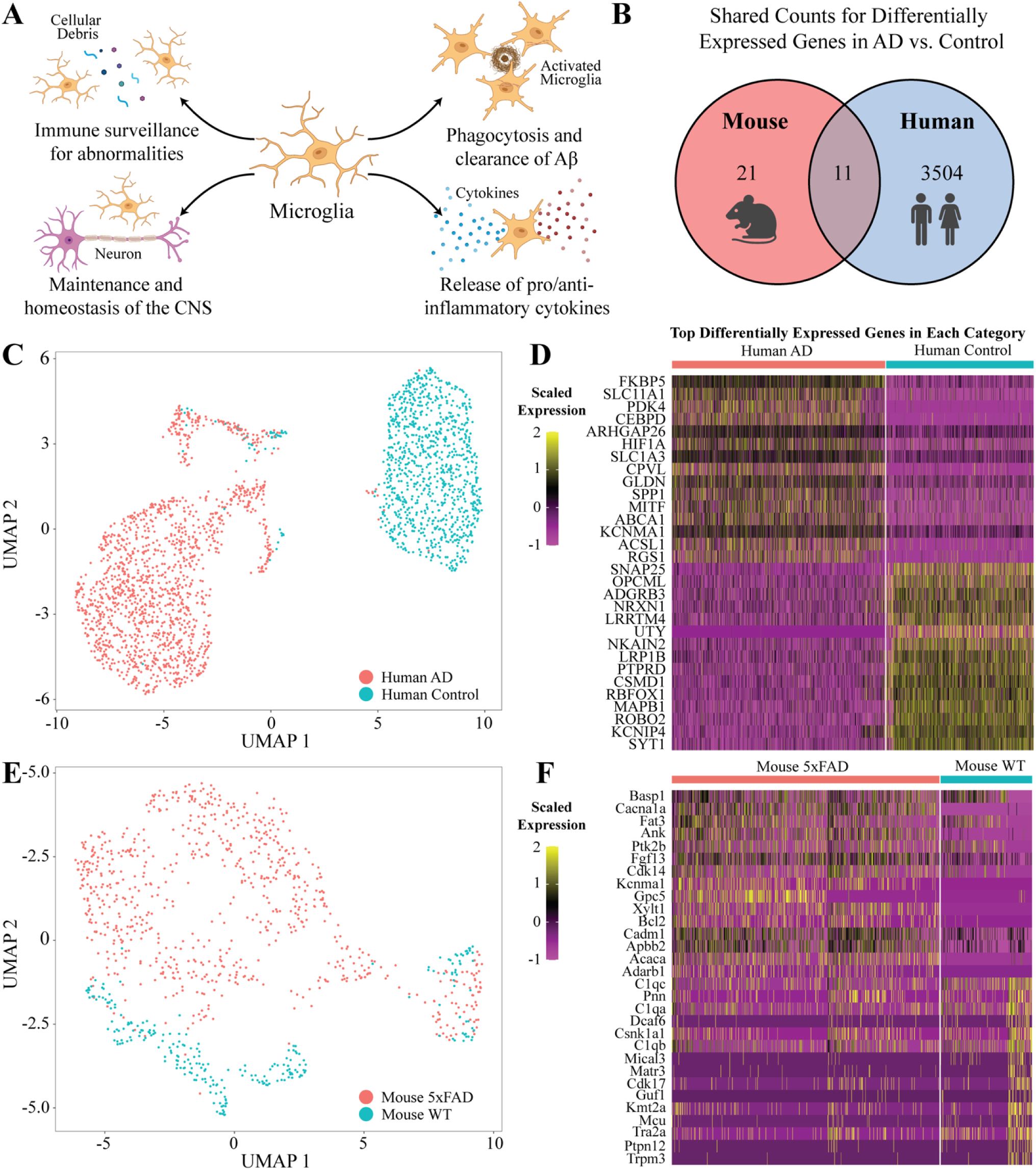
Differential expression analysis of human and mouse microglial cells. **(A)** Microglia mechanism of action in the brain. **(B)** Shared differentially expressed gene orthologs in mouse and human microglial cells. **(C)** UMAP of human single-nuclei RNA-sequencing cells. **(D)** Top 15 differentially expressed genes in human AD (top) and control (bottom) cells. **(E)** UMAP of mouse single-nuclei RNA-sequencing cells. **(F)** Top 15 differentially expressed genes in mouse 5xFAD (top) and wild type (bottom) cells.

We isolated microglia clusters from mouse and human samples for differential expression analysis. The clusters underwent uniform processing to isolate microglial cells further, eliminating possible doublets with marker genes from other cell types. We selected the top 10 variable human microglia PCs to construct the KNN graph with a clustering resolution of 0.5 (**Figure 2C**). We identified 12 clusters, 7 being solely microglia and 5 being cells of other types. After filtering out non-microglial clusters, 2,433 microglial cells (1,438 AD, 995 C) with 1,438 (AD subject 1 = 1,277, AD subject 2 = 161) remain from the AD samples. From the control group, 994 (control subject 1 = 49, control subject 2 = 946) microglial cells remained. Next, we identified the DEGs with an adjusted p-value of less than 0.05 DEGs between the human AD and C samples **(Supplementary Table 3)**. Further, we plotted the top 15 differentially expressed genes in each sample (**Figure 2D)**.

We performed a similar procedure on the mouse microglial cells, where we incorporated the top five variable PCs to construct a KNN graph with a 0.5 clustering resolution **(Figure 2E**). The graph resulted in seven clusters: three composed solely of microglia cells, three of other types of cells, and one with a combination of doublets and microglia cells. The cluster with the combination of doublets was further filtered into smaller clusters to isolate the microglia cells. After removing the non-microglial clusters, we observed 1,319 microglia cells, with 751 coming from the 5xFAD samples (5xFAD mouse 1 = 437, 5xFAD mouse 2 = 314) and 257 coming from the wild-type samples (WT mouse 1 = 190, WT mouse 2 = 67). Next, we identified the DEGs with an adjusted p-value of less than 0.05 between the mouse 5xFAD and WT samples **(Supplementary Table 4)**. The top 15 DEGs in each mouse AD and WT group were also identified (**Figure 2F**). The microglial cells in our mouse 5xFAD model and human AD patients shared a limited number of genes.

### TransComp-R identified mouse principal components that distinguished human AD from control microglial cells

TransComp-R is a computational cross-species translation framework that aims to identify biological signatures in one species predictive of phenotype or therapeutic outcome in another (**Figure 3**). The current implementations of TransComp-R follow the initial step of constructing a principal component space from a model dataset following data normalization and identifying the eigenvectors of the covariance matrix. Further, TransComp-R aims to understand how these eigenvectors explain the variance of a second model dataset. This is done by multiplying these eigenvectors by the normalized data of the second model dataset, ensuring a 1:1 match between genes so we can perform matrix multiplication. Further, we project the second model dataset onto the eigenvectors of the first model dataset, leaving us a PC projection of the second model dataset on the first model dataset. TransComp-R further involves running PC regression to determine if they can significantly predict a phenotype from the second model, validating biological relationships between these models.

**Figure 3.**
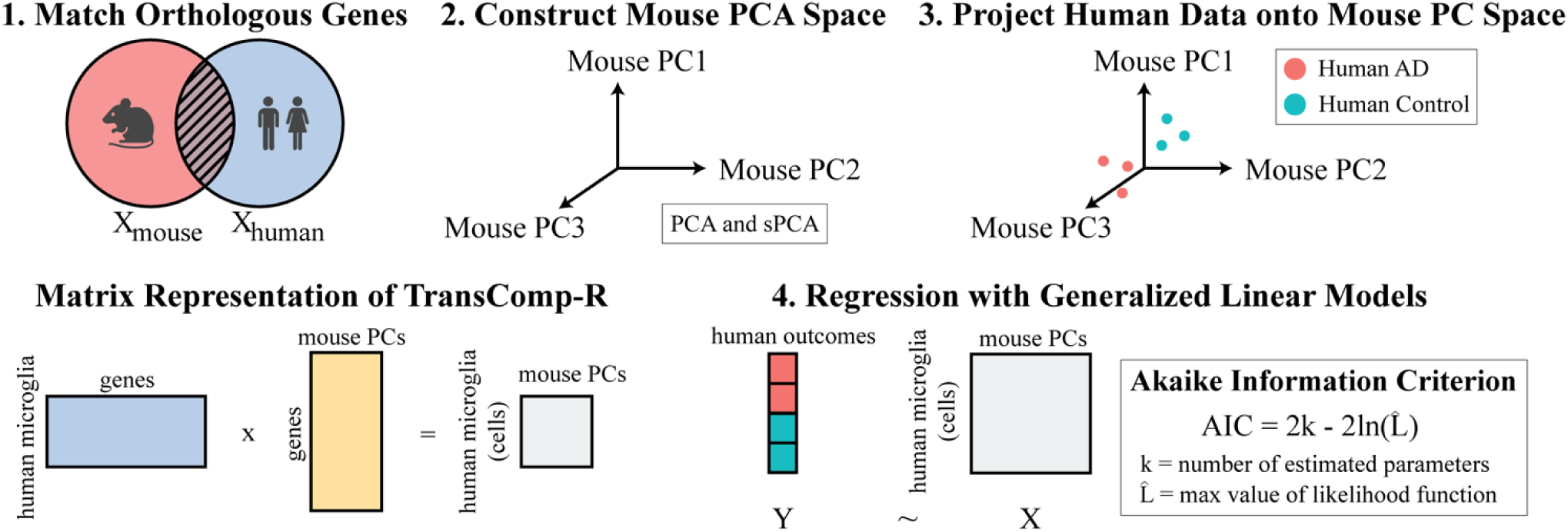
TransComp-R workflow for identification of mouse PCs that significantly distinguish human disease status with single nuclei RNA-sequencing. (1) Mouse and human orthologous genes are matched. (2) Mouse cells leveraging PCA or sPCA techniques establish the mouse PC space. (3) Human cells are projected into mouse PC space by multiplying the human gene expression matrix by mouse PC gene loadings. (4) The resultant human microglia PC projections are regressed against human outcomes to determine which PCs predict human biology.

Our workflow incorporated single-nuclei RNA-sequenced mouse microglia cell data to predict the separation of human microglial cells into control and AD subgroups. While single-nuclei RNA-sequenced data provides vital gene expression information at the cellular level, the datasets become sparser, with more zero-valued measurements (33,34). Compared to previously executed methods of TransComp-R, this model has been tested in limited capacity with sparse datasets (14,18). Thus, we explored the performance of TransComp-R with PCA and sPCA to determine if one approach may be more effective in identifying interpretable and variance-encapsulating PCs across model datasets.

We analyzed the sparsity of our mouse dataset at varying numbers of variable features selected. We found that incorporating different numbers of variable features impacted the relative sparsity of the dataset **(Supplementary Figure S2A)**. Analyzing PCA and sPCA based on AIC **(Supplementary Figure S2B-C)** and AUC **(Supplementary Figure S2D-E)** revealed little to no difference in performance no matter the dataset sparsity. Moving forward, we chose 3,250 variable features to include for TransComp-R as it was the least sparse dataset, containing the most information possible to be captured by PCA and allowing for sparse PCA to introduce as much sparsity as necessary on the genes expressed the most across cells. Next, we generated a mouse PC space and projected human data into the space. The projection of human data into the mouse PC space conserves the direction of the mouse PCs but adjusts the magnitude, thus combining data of the two species. Finally, the human scores projected in the mouse PCs were regressed using generalized linear models and evaluated based on AIC via AIC optimization.

### Sparse PCA and traditional PCA produced similar Results on a technical and biological basis

After implementing TransComp-R and comparing the advantages between sPCA and PCA, we analyzed the PCs to see if we could find biological information relevant to AD. These PCs explained variable levels of variance in mice and humans, with some explaining more variance in humans and some explaining more variance in mice **(Figure 4A-B)**. We noted that PC2, PC3, PC4, and PC5 from regular PCA and sPC3, sPC4, and sPC5 from sPCA explained a high amount of variance in human microglial cells while also having a high level of significance in discerning human AD from control cells **(Supplementary Table 5).**

**Figure 4.**
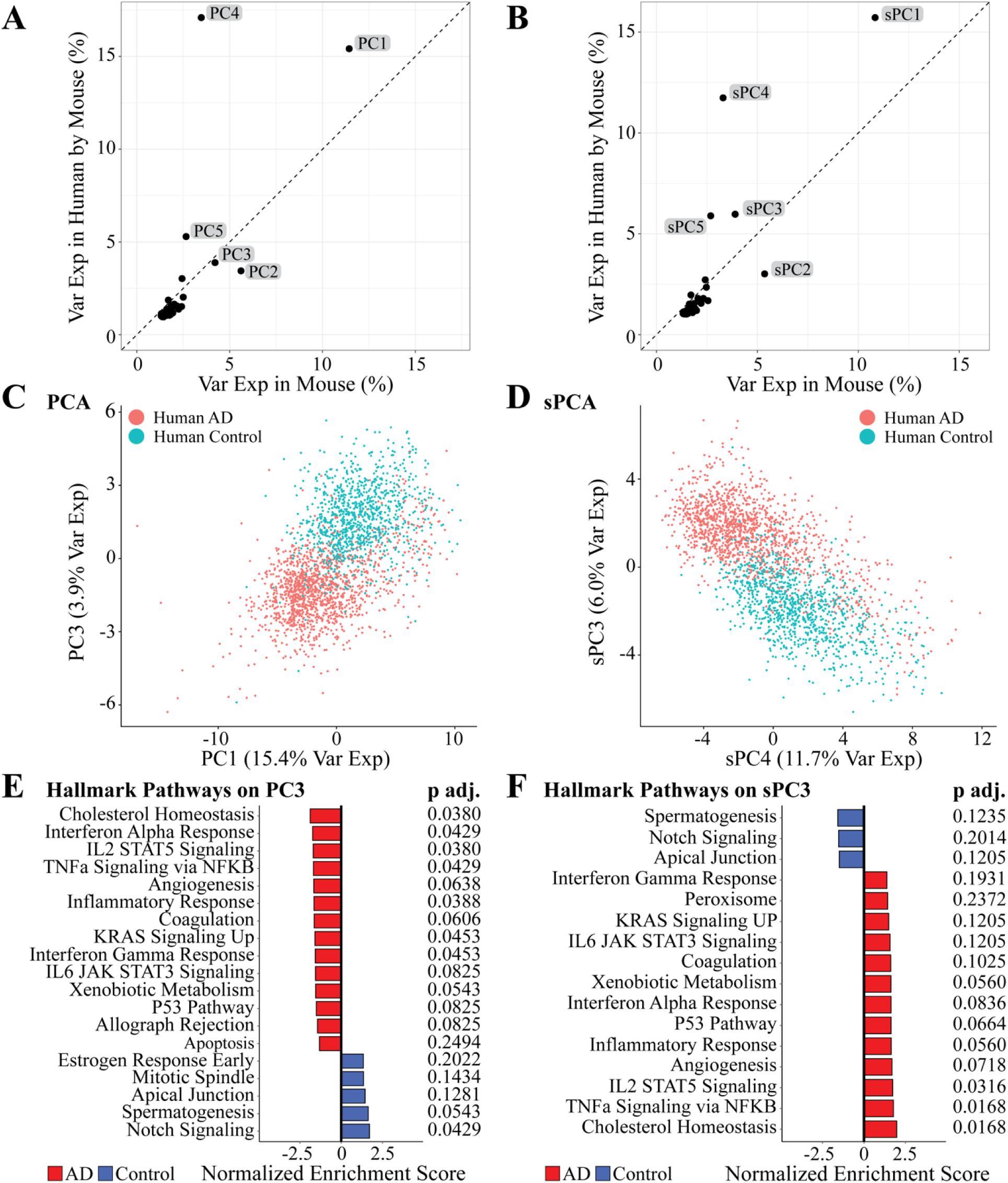
TransComp-R implemented with PCA and sPCA. (A-B) PCs and sPCs variance explained in mouse and human. **(C-D)** Separation of human AD and control groups in both PC3 and sPC3. **(E-F)** GSEA on PC3 and sPC3 using the Hallmark database. Pathway significance was determined with an adjusted *p* < 0.1.

We next performed an AIC optimization to determine which PCs to move forward with for analysis. This identified 22 PCs for PCA and 26 PCs for sPCA to move forward with for downstream analyses **(Supplementary Table 5)**. Looking at regular PCs selected by the AIC optimized model, PC3 was highly significant (*p* < 2e-16) and explained variance in mice (4.21%) as well as humans (3.89%) **(Figure 4C, Supplementary Table 5-6)**. Performing fast gene set enrichment analysis (FGSEA) on the PC loadings, we identified several pathways significantly associated with either AD or control microglial cells **(Figure 4E)**. Specifically, we identified the Cholesterol Homeostasis (*p.adj* = 0.0380), Inflammatory Response (*p.adj*= 0.0429), IL2 STAT5 signaling (*p.adj* = 0.0380), TNFa Signaling via NFKB (*p.adj* = 0.0429), Angiogenesis (*p.adj* = 0.0.0638), Coagulation (*p.adj* = 0.0606), KRAS Signaling Up (*p.adj* = 0.0453), Interferon Gamma Response (*p.adj* = 0.0453), IL6 JAK STAT3 Signaling (*p.adj* = 0.0825), Xenobiotic Metabolism (*p.adj* = 0.0543), P53 Pathway (*p.adj* = 0.0380), and Allograph rejection (*p.adj* = 0.0825) were enriched in the AD microglial cell population. On the other end of the PC, we identified Notch Signaling (*p.adj* = 0.0429) and Spermatogenesis (*p.adj* = 0.0543) as enriched in the microglial cell population.

Transitioning to sparse PCs selected by the AIC optimized model, sPC3 was highly significant (*p* < 2e-16) and explained variance in mice (3.89%) as well as humans (5.97%) **(Figure 4D) (Supplementary Table 5-6)**. Performing FGSEA on the sPC loadings, we were again able to identify several pathways that were significantly associated with either AD or control microglial cells **(Figure 4F) (See Methods)**. Specifically, we identified the Cholesterol Homeostasis (*p.adj* = 0.0168), TNFa Signaling via NFKB (*p.adj* = 0.0168), IL2 STAT5 Signaling (*p.adj* = 0.0316), Angiogenesis (*p.adj* = 0.0718), Inflammatory Response (*p.adj* = 0.0560), P53 Pathway (*p.adj* = 0.0664), Interferon Alpha Response (*p.adj* = 0.0836), and Xenobiotic Metabolism (*p.adj* = 0.0560) were identified in the AD microglial cell population. On the other end of the PC, we identified no significant PCs, although the Spermatogenesis (*p.adj* = 0.1235) and Apical Junction (*p.adj* = 0.1205) pathways had borderline significance.

### Sparse PC4 and PC5 stratified human microglial cells by AD and control conditions

Our AIC-optimized generalized linear model further selected sPC4 and sPC5 to be significant (*p* = 2.95e-12 and *p* < 2e-16), with both sPCs displaying separation between human AD and control groups (**Figure 5A-B, Supplementary Table 5).** sPC4 explained the variance in mice (3.33%) and variance in humans (11.74%), and sPC5 explained the variance in mice (2.69%) and variance in humans (5.89%) **(Supplementary Table 6**). We ran FGSEA for the sPCs to interpret their encoded transcriptomic variance (**Figure 5C-D**). For sPC4, FGSEA identified Hallmark UV Response Downregulation (*p.adj* = 0.0074), Estrogen Response Early (*p.adj* = 0.0871), and Pancreas Beta Cells (*p.adj* = 0.0312) to be enriched in the AD group. On the other hand, Hallmark Oxidative Phosphorylation (*p.adj* = 0.0013) and MYC Targets (*p.adj* = 0.0107) were upregulated in the control. On sPC5, FGSEA identified E2F Targets (*p.adj* = 0.0013) and G2M Targets (*p.adj* = 0.0013) to be upregulated in AD populations with no significant pathways on the control end of the principal component.

**Figure 5.**
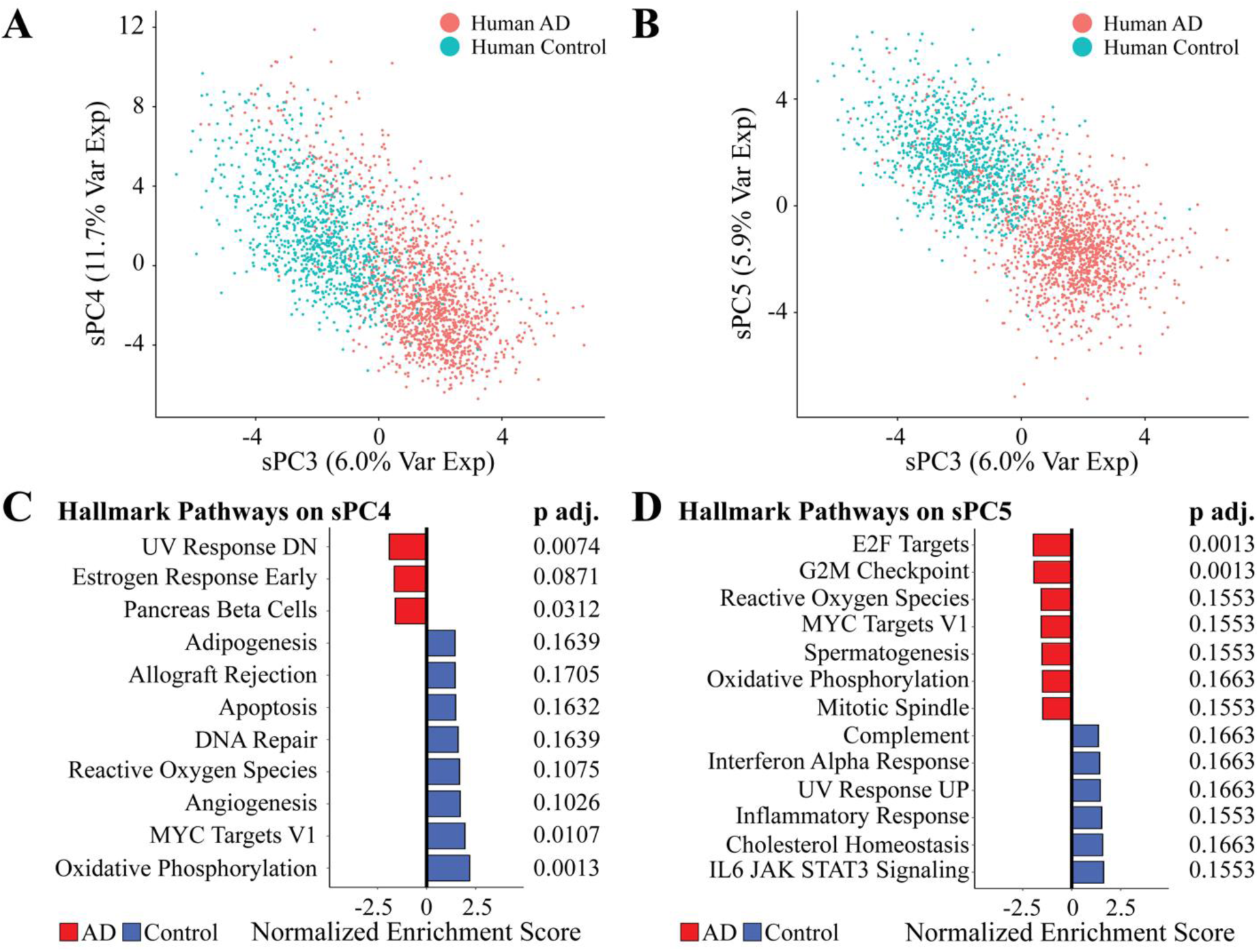
Biological interpretation of the significant sparse principal components. **(A-B)** Separation of human AD and control groups in sPC4 and sPC5. **(C)** FGSEA results of sPC4 **and (D)** FGSEA results of sPC5.

### Computational screening identifies candidate drugs with genetic signature associations to a non-AD Profile

To identify potential drugs correlated with the sPCs predictive of AD and control in microglial cells, we initially filtered from the LINCS L1000 database, 33,609 chemical perturbagens to 2,558 after removing those without gene targets (35,36). The dataset contained characteristic direction coefficients that were used to identify differentially expressed genes, with positive values showcasing upregulation of the gene while negative values showcasing downregulation (37,38). We conducted a Spearman correlation with all the filtered drugs with sPCs that explained a high amount of variance in humans and significantly predicted human AD and control microglial cells, including sPC3, sPC4, and sPC5 based on overlapping differentially expressed genes of the drugs and genes from the sPC’s loadings. We determined 357 significant correlations between drug signatures and sPC3, 416 correlations for sPC4, 202 correlations for sPC5 (*p* < 0.05), and B-H corrected p-values (*q* < 0.05) were highlighted in red **(Figure 6A, C, E)**. After ranking the drugs from lowest to highest correlation values, we emphasized the top three selected drugs sharing similar gene signatures with AD, denoted with (+), while the reverse, denoted as (-) **(Figure 6A, C, E)**. In addition, we determined drugs that had FDA approval and the potential ability to permeate across the BBB (**Figure 6**).

**Figure 6.**
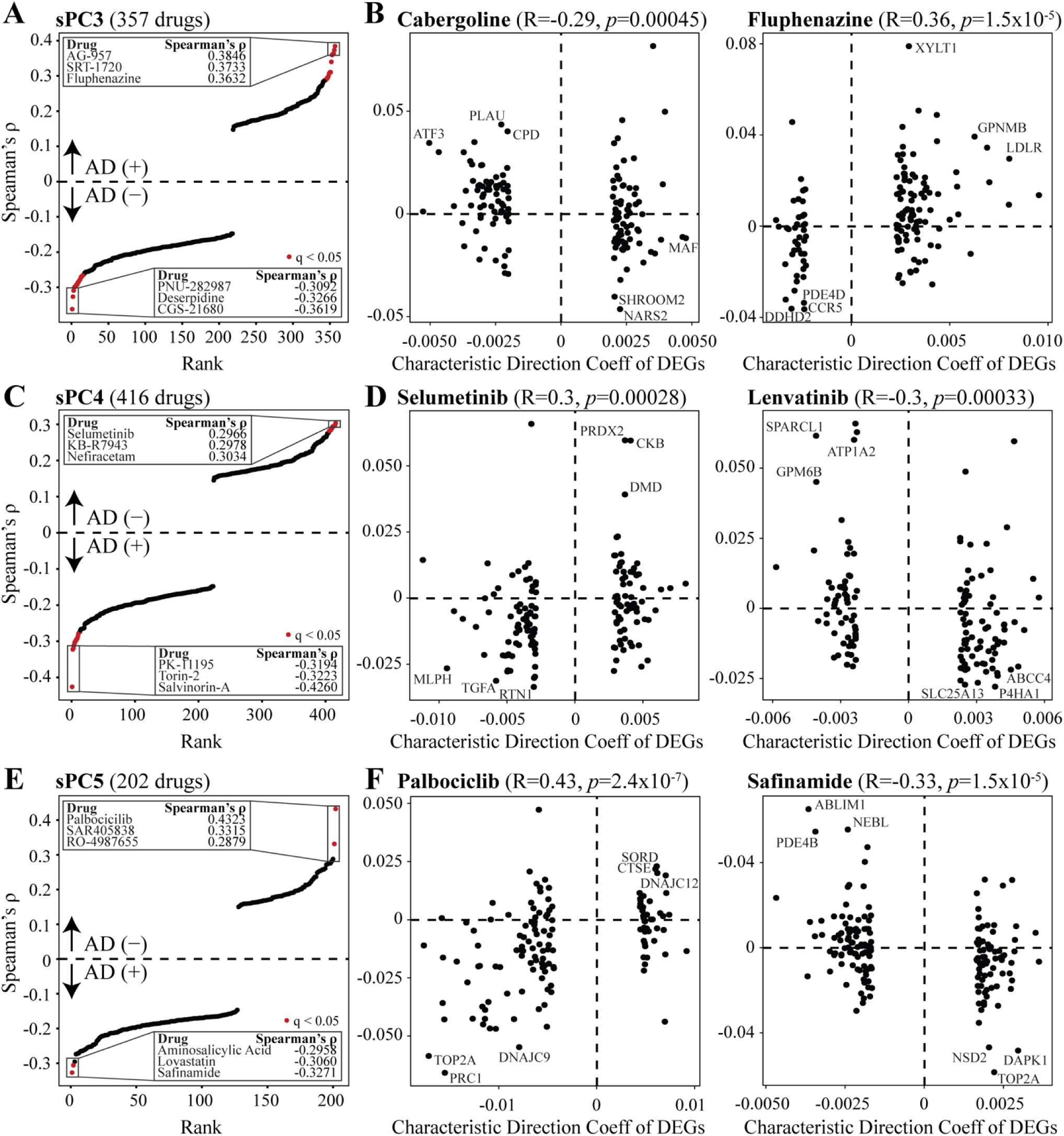
Computational therapeutic screening of drugs. **(A-C)** Significant drugs with Spearman’s correlation values are associated with sPC3, sPC4, and sPC5 genetic signatures. The AD (+) signifies medications related to an AD gene signature, whereas AD (-) signifies medicines associated with a non-AD signature. **(D-F)** FDA-approved medications have the potential to cross the blood-brain barrier.

For sPC3, AG-957 *(ρ*=0.3845*, q*=0.0043), a protein kinase inhibitor (*ABL1*, *EGFR*), SRT-1720 *(ρ*=0.3733*, q*=0.0053), a SIRT activator (*SIRT1*), and Fluphenazine *(ρ*=0.3632*, q*=0.0053), a dopamine receptor antagonist (*DRD1*, *DRD2*), were positively associated with the AD signatures (**Figure 6A**). While the drugs PNU-282987 *(ρ*=-0.3091*, q*=0.0108), a cholinergic receptor agonist (*CHRNA7*), Deserpidine *(ρ*=-0.3266*, q*=0.0053), an angiotensin-converting enzyme inhibitor (*SLC18A2*), and CGS-21680 *(ρ*=-0.3619*, q*=0.0053), an adenosine receptor agonist (*ADORA2A*, *ADORA1*), showed an opposite association with the disease state **(Figure 6A)**. We determined Cabergoline *(ρ*=-0.2918*, q=*0.0425), a dopamine receptor agonist (*ADRA2C*, *DRD3*, *HTR1B*, *HTR2C*, *DRD1*, *ADRA2A*, *HTR2B*, *DRD5*, *DRD2*, *HTR1A*, *HTR2A*, *ADRA2B*, *DRD4*, *HTR1D*, *ADRA1A*), among the negatively correlated drugs with sPC3 AD signature, and Fluphenazine *(ρ*=0.3632*, q*=0.0053), an antipsychotic, with positive correlation with the AD group, to be FDA approved and be able to cross the BBB **(Figure 6B)** (39,40).

We found that drugs on the extreme ends, such as Selumetinib *(ρ*=0.2965*, q*=0.0449), a MEK inhibitor (*MAP2K1*), KB-R7943 *(ρ*=0.2977*, q*=0.0476), a sodium/calcium exchange inhibitor (*SLC8A1*), and Nefiracetam *(ρ*=0.3034*, q*=0.0273), an acetylcholine receptor agonist and GABA receptor agonist (*CHRM1*) to express reverse signatures to the AD group for sPC4 **(Figure 6C)**. In contrast, PK-11195 *(ρ*=-0.3194*, q*=0.0203), a benzodiazepine receptor antagonist (*TSPO*), Torin-2 *(ρ=-0.3223, q*=0.0261), an MTOR inhibitor (*MTOR*), and Salvinorin-A *(ρ*=-0.426*, q*=9.39E-05), an opioid receptor agonist (*OPRK1*) exhibited similar signatures to AD **(Figure 6C)**. FDA-approved drugs that also permeated across the BBB potentially included Selumetinib, used for the treatment of neurofibromatosis type 1 (*NF1*) with positive correlation with the control group, and Lenvatinib, a *KIT*, *PDGFR*, *FGRG*, *and VEGFR* inhibitor (*FGFR1*, *KIT*, *FLT4*, *FLT1*, *KDR*), with a negative correlation with the control group **(Figure 6D)** (41,42).

For sPC5, the chemical perturbagens positively correlated with the control subjects included Palbociclib *(ρ*=0.4323*, q=*6.23E-04), a CDK inhibitor (*CDK4*, *CDK6*), SAR405838 *(ρ*=0.3315*, q*=0.0237), a -, MDM inhibitor (*MDM2*, -), and Ro-4987655 *(ρ*=0.2878*, q*=0.0962), a MEK inhibitor (*MAP2K1*) **(Figure 6E)**. On the contrary, drugs negatively correlated with the control group were Aminosalicylic-acid *(ρ*=-0.2958*, q*=0.1127), a cyclooxygenase inhibitor (*PTGS1*, *PTGS2*), Lovastatin *(ρ*=-0.3059*, q*=0.0492), an HMGCR inhibitor (*HMGCR*), and Safinamide *(ρ*=-0.3270*, q*=0.0191), a monoamine oxidase inhibitor, dopamine uptake inhibitor, and a glutamate inhibitor (*SLC6A3*, *MAOB*) **(Figure 6E)**. FDA-approved drugs included Palbociclib with limited BBB permeability, used for breast cancer, and Safinamide, used for Parkinson’s Disease **(Figure 6F)** (43,44).

## DISCUSSION

The objective of our study was to implement a cross-species computational model to determine murine microglia gene signatures at the single-nuclei level predictive of human AD pathology, and determine potential drugs that inversely regulate AD features. Our processing of mouse and human data and use of known marker genes from literature revealed the same six general cell types: astrocytes, endothelial, microglia, neurons, oligodendrocytes, and oligodendrocyte precursors. This processing allowed us to extract enough mouse and human microglial cells in control and AD populations to make inferences about microglial biology.

A few genes upregulated in human AD that our differential expression analysis revealed were *APOE*, *TREM2*, and *INPP5D* **(Supplementary Table 3)**. GWAS studies have identified these genes as AD risk genes with specific roles in microglial AD pathology, demonstrating relevant biology in these samples (32). A few genes upregulated in the 5xFAD mice that our differential expression analysis revealed were *Cacna1a, Myo1e, Igf1, Apbb2,* and *Gnas* **(Supplementary Table 4)**. These genes have been identified in other 5xFAD models as differentially expressed genes, indicating our mouse samples effectively recapitulate a proven AD model (45,46). When comparing our differential expression analysis of control vs. AD microglial cells in mice and humans, we found that the mouse 5xFAD microglial cells and the human AD samples shared only 34% of differentially expressed genes. This result reiterates that limited biological associations can often be made across AD models, making delineating relevant biological features shared between each system challenging. This fact necessitates using computational methodologies such as TransComp-R to understand these systems’ relationships better.

In our new application of TransComp-R, both PCA and sPCA produced multiple mouse PCs that significantly differentiated human disease outcomes. In evaluating how well PCA and sPCA performed, we found that both on a technical and biological basis, the information encoded on the PCs was similar. At different sparsity levels, there was no clear trend regarding which methodology performed better at a lower or higher sparsity level. For example, using 3,250 genes for our analysis minimized the dataset’s sparsity while including the most significant number of genes; the AIC and AUC for PCA were 740.2127 and 0.981, respectively. For sPCA, the AIC and AUC for PCA were 750.1298 and 0.981, respectively, a minimal difference across the metrics.

Several enriched pathways identified on these regular and sparse PCS were also broadly similar. For example, PC3 and sPC3 shared significant enrichment (*p.adj* < 0.1) of the Cholesterol Homeostasis, Inflammatory Response, IL2 STAT5 signaling, TNFa Signaling via NFKB, Angiogenesis, P53 Pathway, Interferon Alpha Response, and Xenobiotic Metabolism pathways in the AD microglial populations. However, some pathways in PC3 were not significant in sPC3, which can be attributed to specific genes that were removed due to the sparse analysis. This may indicate that sparse principal component analysis may be more stringent and select biological pathways that are likely more relevant. However, we concluded that there is likely no inherent advantage to using one method in the context of TransComp-R, as both effectively captured biological signals in single-nuclei RNA-sequencing data. Although sPCA may offer theoretical advantages in introducing sparsity where genes may not be key drivers in biological pathways—clarifying biological signals (47).

A few of the mouse sparse PCs encoded biology relevant to Alzheimer’s Disease. Looking at how much variance these sparse PCs explained in mice and humans, we found that some explained more variance in humans than in mice, and some explained more variance in mice than in humans. We specifically investigated sPC3, sPC4, and sPC5, which were highly significant in distinguishing human AD from control samples and explained a high amount of variance in humans. We primarily tried to categorize these pathways on relevant PCs into groupings that implicate the biological mechanisms of neurodegeneration, inflammation, and metabolism, hallmarks of AD (48).

Investigating sPC3, we saw an abundance of inflammatory pathways significantly upregulated (*p.adj* < 0.1) in AD microglial samples, including TNFa Signaling via NFKB, IL2 STAT5 Signaling, Inflammatory Response, and Interferon Alpha Response. The literature has shown these pathways to implicate microglia in the brain (49–52). For example, increased TNFa signaling can dysregulate standard microglial phagocytic activity. We also see an abundance of upregulated pathways implicated in neurodegeneration, including Angiogenesis and the p53 Pathway (53,54). The p53 pathway has been shown to increase microglial apoptosis, creating future issues for brain recovery in neurodegenerative diseases such as AD. Finally, we saw the Cholesterol Metabolism and Xenobiotic Metabolism pathways upregulated in AD (55,56).

Turning to sPC4, we saw a few pathways relating to biological mechanisms of neurodegeneration and metabolism. For neurodegeneration, we saw the UV Response Down pathway upregulated in AD and the MYC Targets V1 pathway downregulated in AD microglial cells. Both pathways have literature that supports these findings (57,58). For instance, MYC target genes regulate the early stages of microglial proliferation, indicating a lack of appropriate signaling for proliferation in diseased microglial cells. We also saw several pathways related to metabolism, such as the Estrogen Response Early and Pancreas Beta Cells upregulated in AD, and the Oxidative Phosphorylation pathway downregulated in AD (59–62). A shift from oxidative phosphorylation to glycolysis for energy metabolism in microglia is present in several neurodegenerative diseases, including AD. Finally, looking at sPC5, we saw limited pathways related to neurodegeneration, inflammation, and metabolism. Instead, we saw the E2F Targets and G2M Checkpoint pathways upregulated in AD. As reported in the literature, these pathways are hallmarks of the cell cycle, indicating increased proliferation of microglia in AD in response to brain injury (63).

The finding of altered pathways related to neurodegeneration, inflammation, metabolism, and the cell cycle in AD microglia on these sparse principal components demonstrates that the developed workflow effectively captures relevant disease biology from mice to humans. Additionally, the sparse PCs identified offer a tool to leverage to find drugs that can potentially counteract the dysregulated mechanisms we see by inducing an inverse expression profile of the diseased microglial cells. This will allow for the identification of novel therapeutics for a disease that currently has limited pharmacological treatments available, those of which are available having poor treatment outcomes.

The screening from the LINCS L1000 database revealed drugs with FDA approval and associations with AD and the control group gene signatures. For sPC3, we identified Cabergoline, a dopamine receptor agonist used for hyperprolactinemia and less commonly for Parkinson’s Disease, to showcase opposite signatures to the AD subjects (64,65). Researchers have shown that Cabergoline inhibits extracellular signal-regulated kinase (ERK) signaling from killing neurons by promoting excess glutamate removal (66). In contrast, Fluphenazine, a dopamine receptor antagonist used as a conventional antipsychotic, demonstrated gene signatures in resemblance to the AD group, where it may increase mortality rates among the older population (67). Selumetinib, a MEK inhibitor for treating neurofibromatosis type 1, was positively associated with the control group in sPC4 (68). Acrolein, involved in AD, has been revealed to likely impair the formation of neuron projections and promote oxidative damage. Its harmful effects are counteracted when exposed to Selumetinib (69). Lenvatinib, a multiple tyrosine kinase inhibitor used against cancer, on the other hand, had a negative association with the control subjects with limited studies on the potential efficacy for AD (70).

For sPC5, Palbociclib, a CDK inhibitor used for breast cancer, had a positive correlation with non-AD subjects (71). CDK inhibition using Abemaciclib, another drug with FDA approval, demonstrated decreased amyloid-beta depositions and diminished activated microglia clustering around amyloid-beta aggregates in an AD mouse model (72). However, the inefficient BBB permeability of Palbociclib confers the need for further studies to test the effectiveness of other CDK inhibitors for ameliorating AD symptoms (73). Safinamide, a monoamine oxidase, dopamine uptake, and glutamate inhibitor for Parkinson’s Disease, was negatively associated with the control group (74). Despite exhibiting reverse signatures with the control subjects, Safinamide has restricted the harmful effects of amyloid-beta (75). Our drug screening methodology provides an initial groundwork of potential drugs with associations to microglia mouse gene signatures affected in human AD.

Limitations of our study include limited metadata and demographic information from the publicly available mouse and human datasets due to constraints when selecting matching data across the species. Furthermore, the datasets are restricted to only the nucleus, with no expression values coming from the cytoplasm as both studies performed single-nuclei RNA-sequencing on the brain samples, in which might limit biological signal captured, encapsulating full microglia characteristics (76). In addition, we only explored the 5xFAD mouse model, in which may not capture some features of AD progression in humans. Future studies should investigate different demographic features, including sex, age, and genetic risk factors that could affect microglia in exacerbating AD onset and different mouse models that encompass late-onset AD hallmarks.

Our study provides a new application of TransComp-R by accounting for sparsity, using sparse PCA to build our cross-species integration and taking advantage of single-nuclei RNA-sequencing data to examine specific signatures of microglia in AD. We found biological pathways pertaining to neurodegeneration, inflammation, metabolism, and cell cycle characteristics of mouse microglia that separate AD from healthy controls in human microglia. In addition, we identified drugs from the LINCS L1000 database with potential associations to AD based on the model selected microglia features. Our workflow and results show the effectiveness of our application of TransComp-R for potential drug discovery use.

## MATERIALS AND METHODS

### Selection criteria for publicly available data

We accessed publicly available data through Gene Expression Omnibus (GEO), obtaining single-nuclei RNA sequenced mouse (GSE208683) and human (GSE175814) data (26,27). We selected the mouse and human datasets based on similarities across species, including having the same number of replicates for AD and wild type (WT) samples, sequencing of the same sample type using the same library preparation method (10X Genomics Chromium 3’), and similar cell counts for analysis of approximately 10,000 for control and AD in humans and mice.

The GSE208683 dataset contained single-nuclei RNA brain hemisphere samples from WT, 5xFAD (5x Familiar Alzheimer’s disease), Cnp^-/-^ (Cnp-null mutant), and Cnp^-/-^ 5xFAD (Cnp-null mutant 5x Familiar Alzheimer’s disease) transgenic mouse lines. For our study, we used the WT and 5xFAD samples. There were two wild-type samples with a total of 15,013 cells (WT mouse 1 = 8,212, WT mouse 2 = 6,801) and two 5xFAD samples with 14,662 cells (AD mouse 1 = 7,437, AD mouse 2 = 7,225), which were in the format of pre-filtered matrices with associated metadata. For the human dataset, GSE175814 contained post-mortem Braak stage 3 AD and control samples from the Brodmann area (BA) 41/42, BA6/8, and anterior hippocampal cortex regions of the brain. There were two control samples with 17,214 cells (Control subject 1 = 6,262, Control subject 2 = 10,952) and two AD samples with 11,555 cells (AD subject 1 = 6,223, AD subject 2 = 5,332) before quality control filtering in the format of sparse data matrices from Cell Ranger.

### Data pre-processing and quality control

All computational analysis was conducted with RStudio (ver. 4.1.3). We identified orthologous genes between mouse and human datasets with the *Orthogene* package (ver. 1.0.2) in R (77). We removed duplicate mappings across mice and human species to ensure a one-to-one match before applying the TransComp-R method.

All publicly available data was pre-processed with the *Seurat* R package (ver. 4.3.0). The mouse data was pre-filtered for genes (n < 200) and total transcripts (n < 500). We verified that all mouse mitochondrial gene content was less than 10% for quality control. Additionally, human datasets were loaded and filtered based on mitochondrial gene content and RNA library size **(Supplementary Figure S1A-D)**. We removed samples with a mitochondrial gene percentage greater than 10 or with a median library size less than or greater than three median absolute deviations from the median library size from the analysis. We scored human and mouse cells for regressing cell-cycle genes during data scaling. Finally, each sample, separated by species, was normalized, scaled, and centered by *SCTransform* in *Seurat* with the Gamma-Poisson generalized linear model. We also regressed Cell-cycle scores with *SCTransform*. Samples of the same species were integrated using *Harmony* (ver. 0.1.1) and merged into a singular object by species **(Supplementary Figure S1E-F)** (77).

### Uniform manifold approximation and projection clustering

We performed PCA on the objects used in clustering. A k-nearest neighbors graph was constructed based on the principal components, and the cells were clustered using the Louvain algorithm. Next, we performed a uniform manifold approximation and projection, a dimensional reduction technique with *Seurat* (ver. 4.3.0) to visualize the clustering. Microglia were also re-clustered following the same workflow for doublet removal and data cleaning in mice and humans.

### Differential expression of clusters and disease status

We identified marker genes for clusters in mice and humans during the initial clustering and the microglial re-clustering using the *MAST* (ver. 1.20.0) differential expression test (78). *MAST* is a differential expression package that accounts for common issues in single-cell RNA sequencing, such as stochastic dropout and bimodal expression distributions. The effect of sample-to-sample variability was accounted for by regressing the sample number during DEG identification. The *MAST* package was also employed to identify differentially expressed genes between mouse 5xFAD and WT cells and human AD and control cells.

### Determination of genes to use for PCA and sPCA

We analyzed the sparsity of our mouse dataset at varying numbers of variable features selected via *Seurat* FindVariableFeatures(). We found that incorporating different numbers of variable features impacted the relative sparsity of the dataset **(Supplementary Figure S2A)**. Analyzing PCA and sPCA based on AIC **(Supplementary Figure S2B-C)** and AUC **(Supplementary Figure S2D-E)** revealed little to no correlation between PCA and sPCA based on variable features selected and performance. Thus, 3250 genes were selected for analysis as it minimized the sparsity of the dataset while including the greatest possible number of genes.

### Principal component analysis and sparse principal component analysis

We used the normalized mouse dataset for PCA and sPCA. We first used *Seurat* (ver. 4.3.0) for PCA to scale and center the mouse dataset. Utilizing the top 3250 variable genes identified by Seurat, we then used the top 50 principal components (PCs) for the Translational Components Regression (TransComp-R) framework. For sPCA, we scaled the mouse data using *Seurat* (ver. 4.3.0) and then used the *mixOmics* (ver. 6.18.1) R package to center the mouse dataset and identify 50 PCs for the TransComp-R model.

### Translatable components regression

Our group previously developed Translatable Components Regression (TransComp-R) to overcome interspecies discrepancies between mouse models and human subjects (14). The methodology of TransComp-R projects human samples into mouse PC space. This allows for the variance the mouse model explains to be related to the human data.

We applied TransComp-R to mouse and human single nuclei RNA-sequencing data in this case. We performed regular principal component analysis and sparse principal component analysis on the mouse gene expression matrix to obtain a feature loading matrix with 50 principal components of our mouse data. The mouse feature loading matrix was transposed and multiplied by a human gene expression matrix with orthologous genes. This projected the human cells into the mouse principal component space. The GLM used these PC projections to determine which PCs were significant in discerning human AD from control microglia cells.

### Cross-species variance explained of mouse in human

To determine how well the mouse PCs recapitulate human data, we calculated the percent variance explained by mice in humans. We determined the percentage variance explained with the following formula:

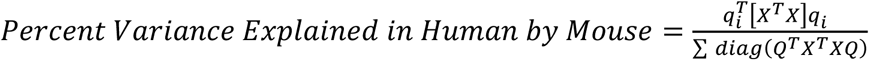

where *Q* is the mouse matrix containing PCs and genes, and *X* is a matrix with human subject information and genes.

### Generalized linear modeling

We created a generalized linear model to determine which PC projections in regular PCA and sPCA had components that effectively separate human AD and control cells. We optimized AIC through the *MASS* (ver. 7.3-60.2) package via the stepAIC function. The following formula calculates the Akaike information criterion (AIC):

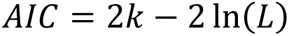

where *k* is the number of estimated parameters in the model, and *L* is the maximum value of the likelihood function for the model. We further analyzed the PCs selected by the AIC optimization for enriched pathways and drug associations.

### Fast gene set enrichment analysis

Pre-ranked FGSEA was employed from the *fgsea* (ver. 1.20.0) package to identify the biological pathways that best described the separation between AD and control groups. We converted mouse genes to their human orthologs and pre-ranked the PC loadings by their respective PC scores for FGSEA. We used the Human MSigDB Hallmark Gene Sets data curation for pathway enrichment analysis and default parameters from the FGSEA package (79,80).

### Computational drug association screening analysis

We obtained the chemical perturbations consensus signatures with characteristic direction coefficients and the small molecules metadata from the Library of Integrated Network-Based Cellular Signatures (35,36). Using the metadata, we filtered chemical perturbagens with established targets and removed duplicates. We found differentially expressed genes for each drug by obtaining the z-scores of their characteristic direction coefficients and selecting genes with p-values < 0.05 for downstream analysis (37,38). We applied a Spearman correlation analysis between each drug and sPCs’ loadings based on overlapping genes to determine associated signatures. For the selected sPCs, the respective Spearman Rho’s with each drug were ranked from lowest to highest, and p-values were adjusted using Benjamini-Hochberg (q < 0.05). We further filtered to find FDA-approved medicines by addressing the Approved Drug Products with Therapeutic Equivalence Evaluations 44^th^ Edition, Appendix A – Product Name Index. In addition, we identified drugs that could cross the blood-brain barrier based on literature analysis.

## CONFLICT OF INTEREST

The authors declare no competing interests.

## AUTHOR CONTRIBUTIONS

AB: Conceptualization, data curation, formal analysis, investigation, methodology, software, validation, writing - original draft, writing - review and editing. JHP: Conceptualization, data curation, formal analysis, methodology, software, validation, writing - original draft, writing - review and editing. BKB: Formal analysis, software, validation, visualization, writing - original draft, writing - review and editing. DKB: Conceptualization, funding acquisition, methodology, project administration, resources, supervision, writing - review and editing. All authors contributed to the article and approved the submitted version.

## FUNDING

This work is supported by the Good Ventures Foundation and Open Philanthropy, as well as start-up funds from the Weldon School of Biomedical Engineering at Purdue University and Case Western Reserve University (AB, JHP, BKB, DKB). BKB is supported by the National Science Foundation Graduate Research Fellowship Program (GRFP) under grant number DGE-1842166. BKB acknowledges the support of the NIH T32 predoctoral fellowship T32DK101001 from the National Institute of Diabetes and Digestive and Kidney Diseases.

### ACKNOWLEDGMENTS

The authors acknowledge content from BioRender.com used in **Figure 1A**, **Figure 2A-B**, and **Figure 3**.

## SUPPORTING INFORMATION

**Supplementary Figure S1.**
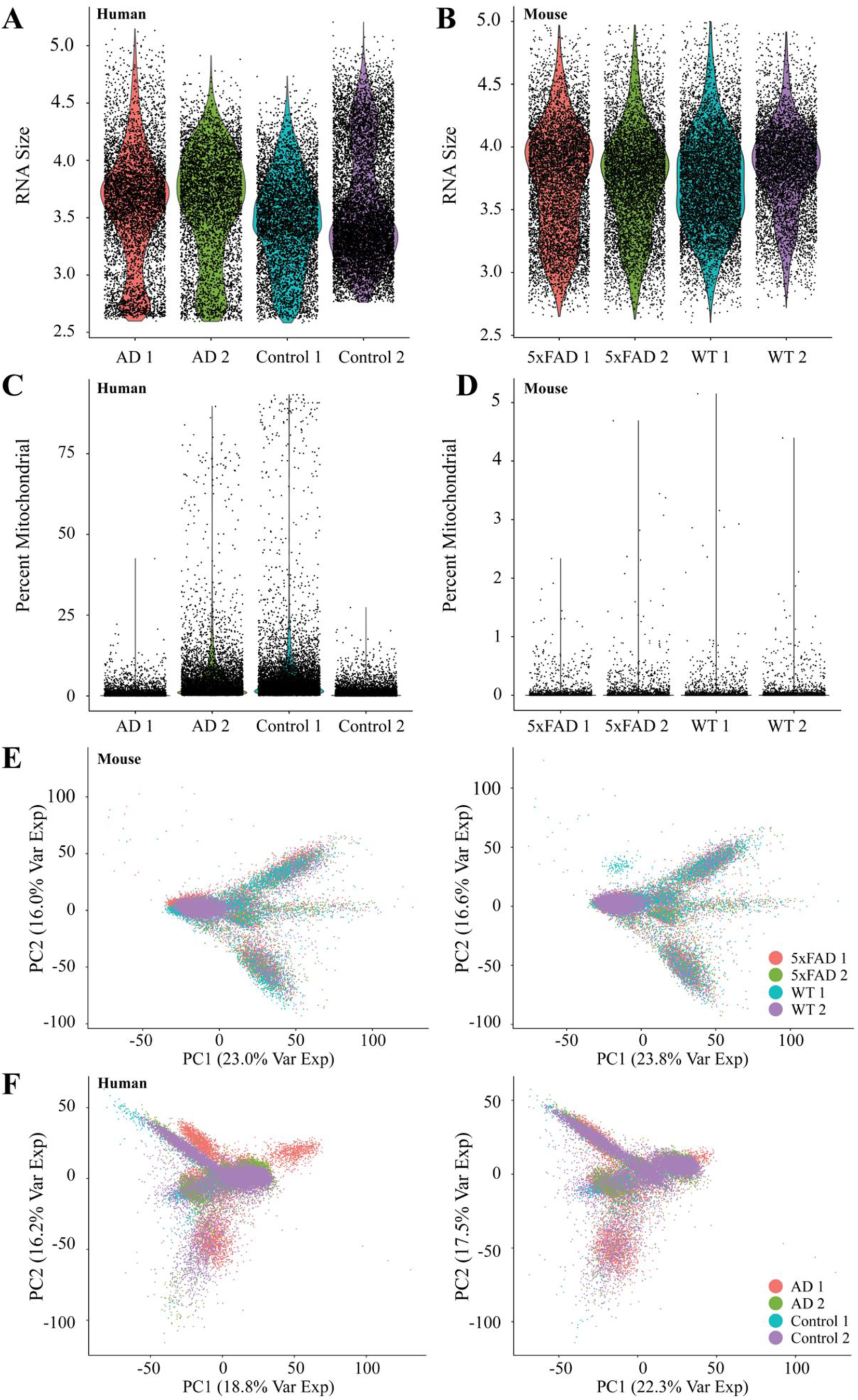
Quality control of human and mouse single nuclei RNA-sequencing data. **(A)** Gene counts of human samples before QC filtering **(B)** Mitochondrial gene content of human samples before QC filtering **(C)** Gene count of mouse samples **(D)** Mitochondrial gene content of mouse samples **(E)** PCA plot of top two principal components in humans before harmony batch correction (left) and after harmony batch correction (right) **(F)** PCA plot of the top 2 principal components in mice before harmony batch correction (left) and after harmony batch correction (right)

**Supplementary Figure S2.**
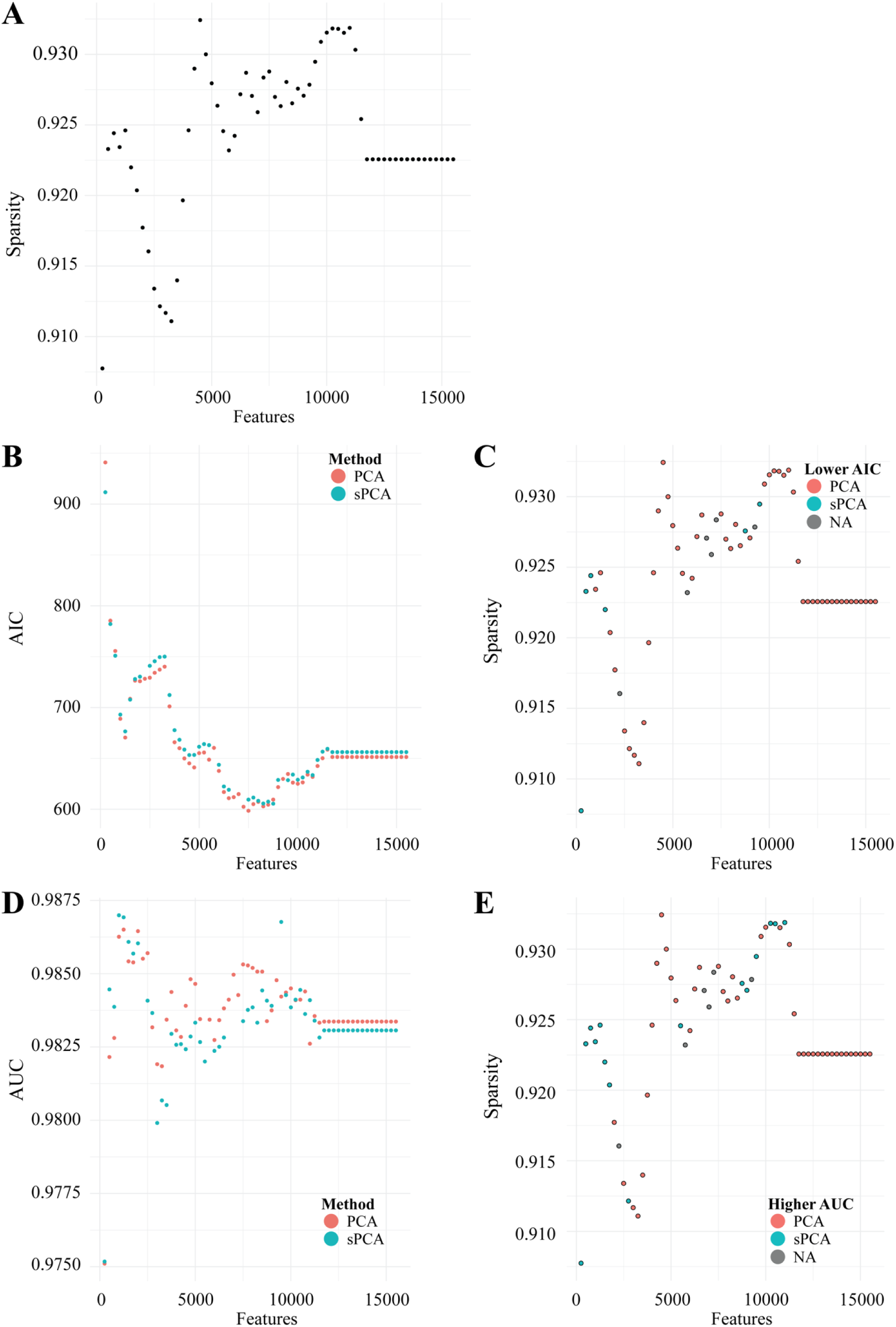
**(A)** Sparsity of data frame as a function of variable features selected **(B)** AIC score as achieved through AIC minimization for PCA and sPCA as a function of variable features **(C)** Sparsity of data frame a function of variable features selected colored by lowest AIC **(D)** AUC score achieved through the ability of AIC minimized PCs to predict disease status as a function of variable features **(E)** Sparsity of data frame as a function of variable features selected colored by highest AUC

## Notes

### Competing Interest Statement

The authors have declared no competing interest.

